# Structural and quantum chemical basis for OCP-mediated quenching of phycobilisomes

**DOI:** 10.1101/2023.09.30.560311

**Authors:** Paul V. Sauer, Lorenzo Cupellini, Markus Sutter, Mattia Bondanza, María Agustina Domínguez Martin, Henning Kirst, David Bína, Adrian Fujiet Koh, Abhay Kotecha, Basil J Greber, Eva Nogales, Tomáš Polívka, Benedetta Mennucci, Cheryl A. Kerfeld

## Abstract

Cyanobacteria employ large antenna complexes called phycobilisomes (PBS) for light harvesting. However, intense light triggers non-photochemical quenching, where the Orange Carotenoid Protein (OCP) binds to PBS, dissipating excess energy as heat. The mechanism of efficiently transferring energy from phycocyanobilins in PBS to canthaxanthin in OCP remains insufficiently understood. Using advanced cryogenic-electron microscopy, we unveiled the OCP-PBS complex structure at 1.6-2.1 Å resolution, showcasing its inherent flexibility. Employing multiscale quantum chemistry, we disclosed the quenching mechanism. Identifying key protein residues, we clarified how canthaxanthin’s transition dipole moment in its lowest-energy dark state becomes large enough for efficient energy transfer from phycocyanobilins. Our energy transfer model offers a detailed understanding of the atomic determinants of light harvesting regulation and antenna architecture in cyanobacteria.

**One sentence summary:** High-resolution cryo-EM structure of the OCP-PBS complex reveals intrinsic motions and enables the atomic simulation of the quenching mechanism

## Introduction

In photosynthetic organisms, light-harvesting antennae capture and direct energy through elaborate networks of protein-scaffolded pigments to ultimately be used to fix CO_2_. In cyanobacteria, the principal antenna complex is the phycobilisome (PBS). In these massive macromolecular assemblies, light-absorbing phycocyanobilin (PCB) pigments are covalently bound to phycobiliproteins that are tethered together by linker proteins and organized into the phycobilisome rods and core cylinders. The protein environment of the pigments tunes their spectral properties allowing for efficient and fast transfer of the excitation energy towards the photosynthetic reaction centers.

The efficiency of photosynthesis relies on balancing the delivery of excitation energy and the ability to photoprotect under conditions in which the harvested light energy exceeds photosynthetic capacity. Non-photochemical quenching (NPQ) is the process by which excess captured light energy is dissipated as heat (*1*). In cyanobacteria, NPQ is carried out by the Orange Carotenoid Protein (OCP), a 34 kDa protein containing a single keto-carotenoid (*2, 3*). It converts from a resting, orange form (OCP^O^) to a red, active form (OCP^R^) after absorption of intense blue light. Two OCP^R^ homodimers then bind to the PBS core, activating quenching (*4*). Although it is indisputable that OCP causes quenching in the PBS, the quenching mechanism itself is still debated (*2*). Excitation energy transfer (EET) from the PBS phycocyanobilin to the OCP-bound keto-carotenoid appears as a promising explanation. Our recent structure of the OCP-PBS complex (*4*) supports this picture, as the keto-carotenoid (canthaxanthin) is located close to two ApcA PCBs. However, EET is only efficient if there is resonance between the donor (PCB) emission and acceptor (canthaxanthin) absorption, and if there is sizable electronic coupling between the two electronic transitions. The first singlet excited state (S_1_) of carotenoids has the right energy to accept EET from PCBs, but it is notoriously optically dark, which makes the electronic coupling small. Conversely, the second excited state S_2_ is optically bright, but its energy is too high to allow EET. A somewhat large transition dipole moment (TDM) for the S_1_ state has to be assumed to match the quenching rates observed experimentally. However, it is not clear how this large TDM is achieved in OCP-bound canthaxanthin.

New developments in cryo-EM data acquisition and processing have made it possible to obtain structures of well-behaved test specimens at near-atomic resolution (≤2 Å), with the promise to extend this resolution goal to more challenging targets in physiologically relevant states (*5, 6*). More accurate atomic models obtained from such high-resolution structures are bringing new insight into the role of local intrinsic protein motions in large macromolecular assemblies and are useful to determine properties of quantum mechanical processes such as pigment excitation that is especially relevant in photosynthetic protein complexes (*7, 8*).

We were previously successful in obtaining cryo-EM structures of quenched and unquenched PBS from the model cyanobacterium *Synechocystis* sp. PCC 6803, but our efforts stepped short of offering a detailed model of energy transfer that considers the local protein environment and positions of associated water molecules (*4*). Such a model is necessary to understand in detail how a small protein with a single carotenoid can fully quench the 6.2 MDa PBS.

Here, we take advantage of the latest cryo-EM developments to determine the structure of the quenched PBS bound to OCP to a resolution of 2.1 Å in the core and 1.8 Å in the rods, with some regions reaching 1.6 Å resolution. This enabled building of an unprecedentedly accurate model of the structure, including ordered water molecules and hydrogen atoms, and the characterization of local intrinsic motions within the PBS. We then used this atomic model to perform multiscale quantum chemical calculations that reveal the interplay between pigments and protein residues enabling OCP to serve as an energy sink for the PBS. We identify several residues within OCP that are critical to establish the sizable transition dipole moment along canthaxanthin that is required for quenching. The energy transfer model derived from our structure explains the fast excitation energy transfer from the ApcA PCBs to canthaxanthin, and provides a better understanding of the atomic determinants of light harvesting and energy transfer and its regulation, the latter including revealing macromolecular motions within the PBS.

## Results

### High-resolution cryo-EM structure of OCP-PBS

To obtain a more detailed understanding of OCP-mediated PBS quenching, we sought to determine a high-resolution cryo-EM structure of the OCP-PBS complex, taking advantage of recent technological developments in data acquisition and processing. We used our previously established protocol for streptavidin affinity grids to stabilize the sample on the cryo-EM grid (*9, 10*) and then acquired the data on a 300 kV cryo-electron microscope equipped with a Falcon 4 direct electron detector with increased DQE at Nyquist frequency, a cold field emission gun (cold-FEG) and an energy filter capable of maintaining a narrow slit width for extended periods of time (*5, 11*). This setup allowed us to maximize the signal-to-noise ratio as compared to conventional microscopes, particularly at resolutions beyond 2 Å. To further increase the resolution and to combat loss of data quality caused by protein complex flexibility intrinsic to all biological macromolecules, we performed 3D variability and 3D flexibility analysis as implemented in recent software packages (*12, 13*). Using these approaches we were able to obtain a structure of the OCP-PBS complex from *Synechocystis* sp. PCC 6803 with a resolution of up to 2 Å in the core and 1.8 Å in the rods (see Methods in the Supplementary Materials), an improvement of approx. 0.3-0.5 Å for the rods and the core, respectively (Fig. 1a,b; Fig S1,2, Table S1). The newly obtained reconstruction recapitulates the known OCP-PBS structure, consisting of a tricylindrical core composed of Apc proteins with six emanating rods composed of Cpc proteins. On each side of the core, one OCP dimer is wedged between the top and bottom cylinders, as previously reported. Within one OCP dimer, the two OCPs connect via their C-terminal domain while their N-terminal domains and their embedded canthaxanthin molecules bind to AcpA and ApcB of the PBS core. The gain in resolution allowed us to build an atomic model of the OCP-PBS with increased accuracy compared to our previous structure (Fig. 1b). High resolution details were readily visible in several parts of the map, for example in the rods, where characteristic holes in the density of aromatic residues became apparent (Fig. 1c). Within the N-terminal domain of the OCP the local resolution of canthaxanthin and its surrounding residues has increased to 1.7 - 2 Å, and therefore the density for all four methyl moieties branching off the polyene chain and for the two terminal beta-ionone rings can be clearly assigned (Fig. 1d,e). The new map also shows more continuous density for the C-terminal domain (CTD) of OCP and the quality of the map for the CTD increased further after accounting for protein flexibility. Regions of OCP-CTD that were previously poorly resolved, especially around the CTD dimerization interface, now show clear continuity and side chain density, confirming the proposed interface (Fig. 1f).

**Figure 1:**
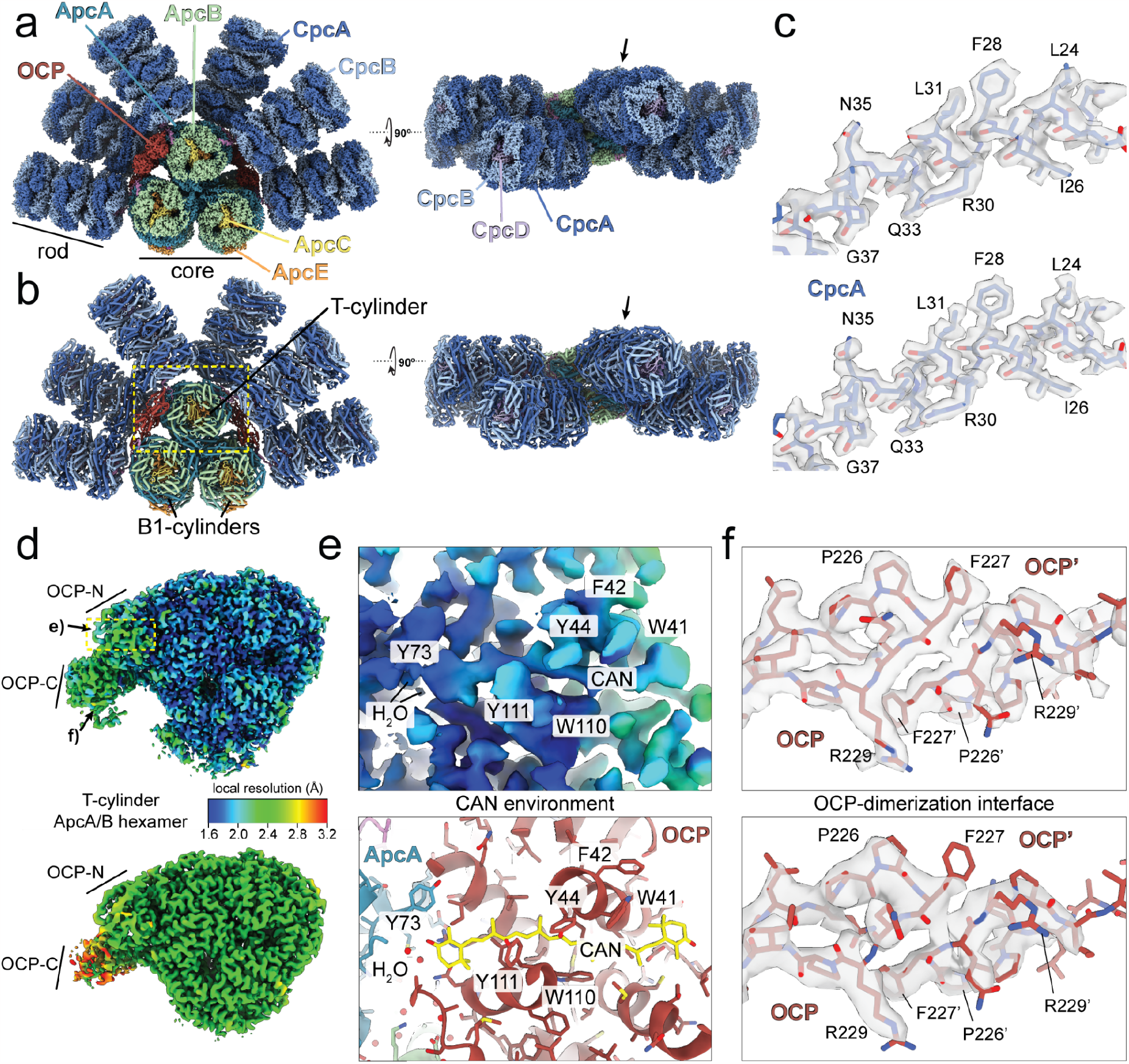
Structure of the OCP-PBS. (a) Composite cryo-EM map of the OCP-PBS colored by subunit type. Arrow points to molecular detail shown in (c). (b) Cartoon representation of the OCP-PBS atomic model with the arrow pointing to molecular detail shown in (c). Yellow dashed box marks the region of the OCP-PBS model shown in (d). (c) Map details for the rod in previous (top) and new (bottom) cryo-EM reconstructions. (d) Local resolution map of the T-cylinder disk bound to OCP in our new reconstruction (top) compared to our previous reconstruction (bottom). Arrows indicate the location of details shown in (e) and (f). (e) Local resolution within the N-terminal domain of OCP surrounding the canthaxanthin (CAN) (top) and corresponding atomic model (bottom). (f) Map quality of the OCP dimerization interface of our previous reconstruction (top) compared to this study (bottom).

In addition, we were able to model more than 7,000 unique water molecules that were previously not visible, including 49 water molecules at the OCP-PBS interface (Fig. 1e), allowing us to include them into our energy transfer and quenching calculations (see below). Additionally, we can identify density for the post-translational modification of residue Asn72 of CpcB to N4-methyl-asparagine (Fig. S3) (*14, 15*).

Analysis of the data using ResLog indicated that resolution of our maps was limited mainly by particle numbers and not by sample heterogeneity (Fig. S4a) (*16*). Therefore, to explore to which extent resolution detail can be improved, we took advantage of the local D3 symmetry within the central Cpc(*αβ*)_6_ double hexamer of the rod (“central rod disk”) to improve the signal-to-noise ratio. Applying D3 symmetry resulted in a local map of the central rod disk with a resolution of 1.6 Å, with the details expected at this resolution clearly visible (Fig. S4b-d). The same particles yielded a resolution of 1.8 Å when not applying any symmetry (C1). Calculating difference maps allowed us to visualize hydrogen density along the c-alpha backbone of several alpha helices (Fig. S4d). We believe that the use of streptavidin affinity grids was particularly critical to get significantly higher resolution than any other published PBS structure so far. This type of grid greatly alleviated the issue of preferential orientation of the PBS and protected the particles from harmful interactions with non-biological interfaces, such the air-water interface.

Previous cryo-EM reconstructions of the cyanobacterial phycobilisome suffered from inherent flexibility of the complex that made rods appear ‘smeared out’ or incomplete (*4*). The high quality of our data and recent software developments allowed us to both increase the overall map quality of several domains of the PBS (Fig. 1c-e), and to extract biologically significant information about conformational variability. Using 3DVA and 3DFlex (*12, 13*) we investigated the intrinsic movement on three scales: the entire OCP-PBS, its tri-cylindrical core, and the T-cylinder bound to OCP (Fig. 2). The overall movement of the holo-OCP-PBS complex is dominated by movement of the 6 rods that emanate from the core region (Fig. 2a, Movie S1). The rods rock ‘up’ and ‘down’ in small angles as rigid units relative to the core. The overall displacements are in the range of 20 Å and they are less pronounced in the ‘forth’ and ‘back’ directions. We previously proposed that the rods in the PBS are mobile and that they can adopt distinct conformations that control access of OCP to the PBS core (*4*). Our flexibility analysis shows that the rods have an inherent propensity to move, in agreement with that hypothesis. We anticipate that within the cell the intrinsic mobility will have an impact on the supramolecular assembly and disassembly of PBS arrays to regulate light harvesting.

**Figure 2:**
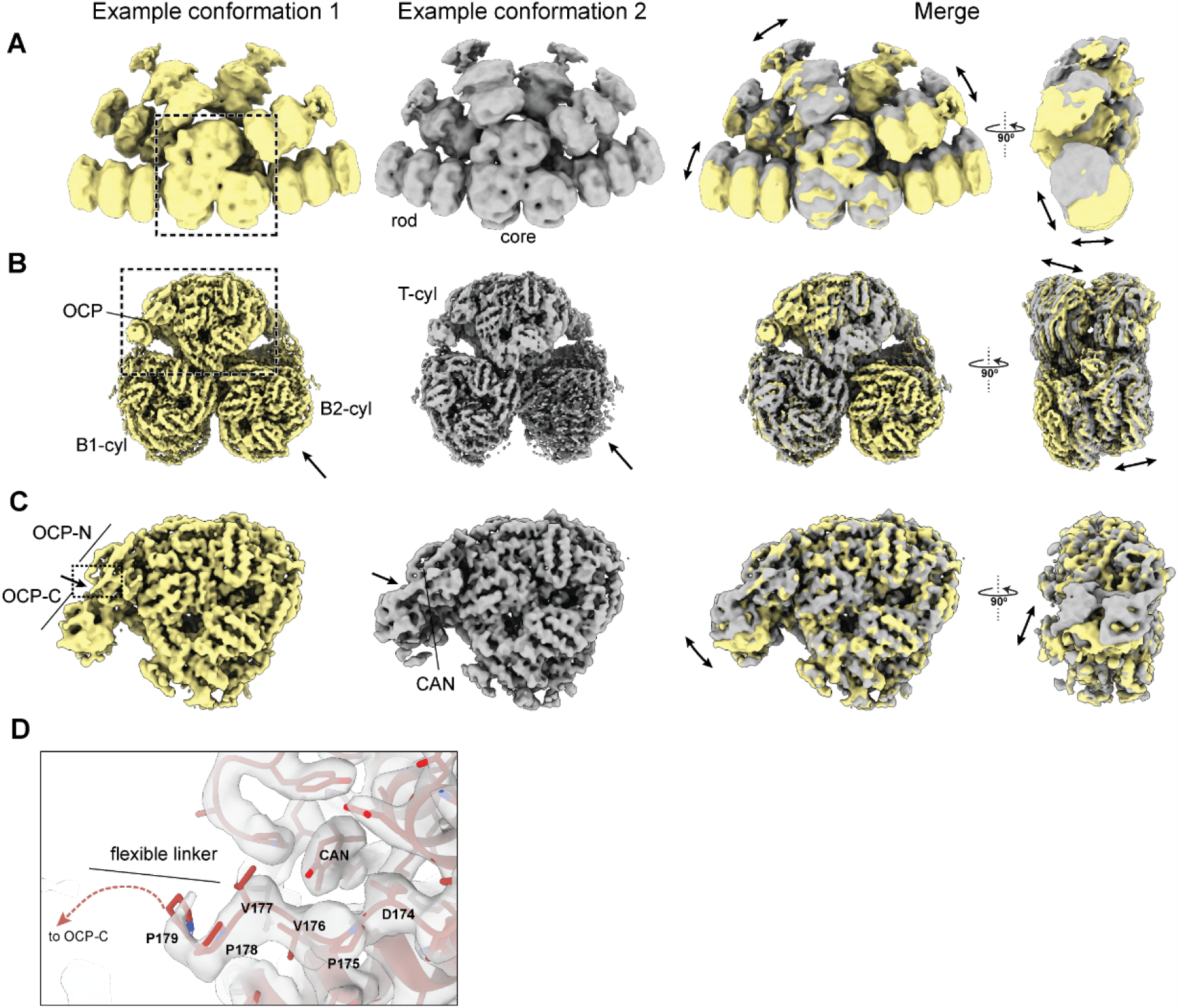
Visualization of distinct forms of motion in OCP-PBS at different scales (A)-(C). The two extreme states within a certain motion are displayed next to each other (yellow and gray) and the superimposed and displayed in two orthogonal views. (A) Holo-OCP-PBS. The double arrows indicate the direction of rod movement. (B) OCP-PBS core. The arrow points to the position of loss of ApcA/B double hexamer in the B2 cylinder and the double arrows indicate the direction of twisting movement. (C) T-cylinder disk bound to OCP. The single arrow points to the location of the flexible linker between the N- and C-terminal domains of OCP and the double arrows show the direction of OCP-CTD movement. The dashed box shows the part of the structure that is enlarged in (D). (D) Molecular details around the flexible linker of OCP. The linker residues proximal to the CAN molecule are indicated.

The core of the OCP-PBS complex displays an overall slight twisting motion, with the top cylinder and bottom cylinders moving in opposite directions around the central symmetry axis (Fig. 2b, Movie S2). Our analysis also revealed that one of the B-cylinder’s ApcA/B hexamers is lost in a subset of particles. While this could occur during cryo-EM sample preparation, it agrees with previously reported sample instability or heterogeneity and may give rise to the presence of spectroscopically active components in PBS samples that are distinct from the holo- complex (*17, 18*).

Within the isolated T-cylinder disk bound to OCP structural flexibility appears mostly restricted to the motion of the dimerized CTDs of OCP with respect to the remainder of the complex (Fig. 2c, Movie S3). The most prominent motion of the dimerized CTDs of OCP appears to be an ‘up’ and ‘down’ rocking relative to the core. The flexible linker connecting the NTD and CTD of OCP moves in concert with the CTD dimer. Residues D174-V176 within the linker are in close proximity to the Canthaxanthin (CAN) molecule that is embedded in the NTD of OCP^R^ (Fig. 2d). Because the protein environment of CAN impacts its electronic properties and therefore its ability to perform NPQ, we asked whether the intrinsic flexibility of OCP would play a role in regulating this effect. To answer this question we used our new atomic model of the OCP-PBS to calculate the excited state parameters of CAN and to determine the mechanism of CAN mediated NPQ in detail.

### Multiscale quantum chemical calculations on OCP-PBS

Our previous model describing the energy transfer quenching of PBS by OCP (*4*) used empirical values for two key parameters of the carotenoid: the transition dipole moment (TDM) of canthaxanthin in OCP and its S_0_-S_1_ transition energy. Here, in contrast, we exploit our higher-resolution structure to apply quantum mechanics/molecular mechanics (QM/MM) calculations in addition to restrained molecular dynamics (restMD) simulations to obtain more realistic values of these parameters accounting for the effects of CAN-protein interactions (see Methods in the Supplementary Materials). Given the large size of the entire PBS core, in our OCP-PBS model we considered only the closest regions of the core to the OCP-NTD, which is bound to the T-cylinder of PBS. To sample solvent configurations, we performed restMD simulations in explicit water solvent where the system was restrained on the backbone of the proteins of our high-confidence model. Sampling of water configurations is essential to accurately determine the effects of the solvent on the electronic structure of CAN. In fact, it has been observed that bulk water can partially screen the protein’s electrostatics in both OCP^O^ and NTD (*19*). As evident from cryo-EM, the OCP-CTD and its linker region are inherently flexible, which could tune the electronic properties of CAN. To investigate this hypothesis, the linker was left unrestrained in the restMD simulations to fully sample its configurational space. Due to the large distance between CAN and the C-terminal domain, the latter was excluded from the model. In agreement with the structural data, our restMD simulations show that the linker exhibits substantial flexibility (Figure 3a), and that it can acquire different conformations and positions. Furthermore, the regions of high water occupancy in the restMD simulations mirror the positions of resolved water molecules in the cryo-EM structure (Figure S6) and the positions of CAN and the closest polar residues sampled by restMD fit well into the cryo-EM map (Figure S9).

**Figure 3:**
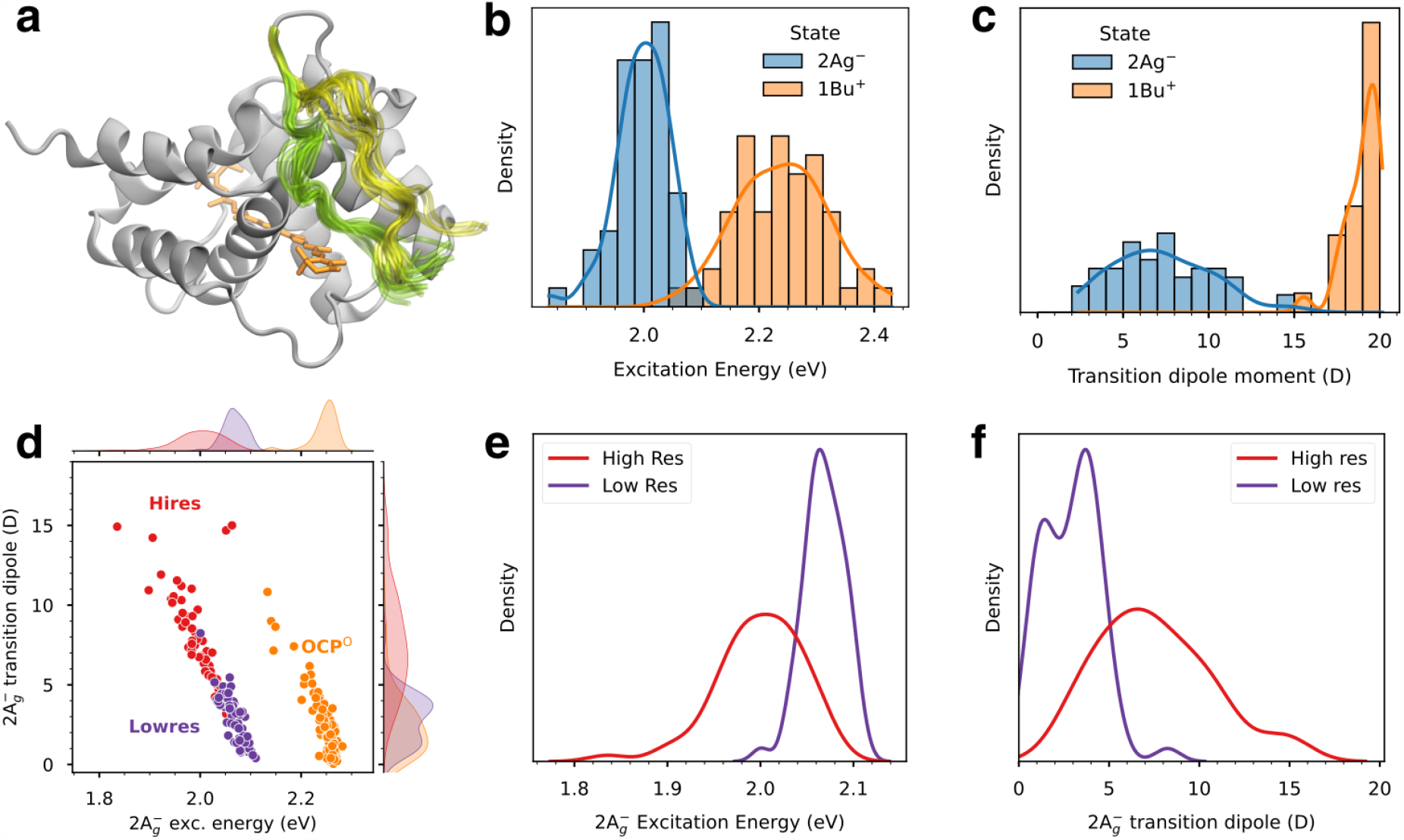
Linker flexibility and CAN excited-state properties. (**a**) Flexibility of the linker in our restMD simulations. Representation of the NTD in OCP-PBS (atoms of the PBS are omitted) along two restMD replicas. The colored parts (yellow and green for the two replicas) correspond to the linker residues G169–P179 that were allowed to move during the MD. CAN is shown in orange. (**b**,**c**) Distribution of the excitation energy (**b**) and transition dipole moment (**c**), for the dark (2A_g_^-^) state and the bright (1B_u_^+^) state calculated along the restMD trajectories. (**d**) Comparison of dark-state properties in OCP-PBS and OCP^O^. Points represent pairs of excitation energy and TDM for each structure sampled from OCP^O^ (orange) or OCP-PBS (red), whereas the shaded curves on top and right side represent the marginal distribution of excitation energy (top) and TDM (right). Points corresponding to the previous OCP-PBS structure (*4*) are shown in purple. (**e**,**f**) Comparison between CAN S_1_ properties obtained with QM/MM calculations on restMD simulations from higher resolution structure in this study and that for our previously published structure. (**e**) Excitation energy of the S_1_ state, (**f**) TDM of the S_0_-S_1_ transition.

#### Excited-state properties of CAN

Traditionally, the photophysics of carotenoids is described by a three-level model, the ground state (S_0_) and the two lowest excited state (S_1_ and S_2_). Moreover, the S_0_-S_1_ transition (generally indicated as 2A^−^_*g*_) is optically forbidden (dark) and the light is absorbed by the S_2_(1B_u_) bright state. Following this model, we extracted structures for the OCP-PBS model from two independent restMD trajectories, and calculated the transition energies and the corresponding TDMs for the S_0_-S_1_ and S_0_-S_2_ transitions of CAN using a semiempirical QM method (*20*), coupled to a molecular mechanics (MM) description of the protein matrix and the solvent. We find that, in agreement with earlier hypotheses (*4*), the S_0_-S_1_ TDM of CAN in OCP-PBS is substantially increased with respect to isolated CAN (Table 1), although it remains always significantly lower than the one found for the S_0_-S_2_ bright transition (Figures 3b,c). Notably, we observe a broad distribution of TDM values for the dark state, sometimes exceeding 10 D, and with an average (7 D) much larger than the 2.3 D obtained from previous calculations on echinenone in OCP^O^ (*21*). As in our restMD, the well-resolved regions of the protein are restrained, together with the CAN, the TDM distribution only arises from the dynamics of water and external side chains, which modulate the protein electric field acting on the embedded carotenoid.

**Table 1:** Excitation energies (Δ*E* in eV) and transition dipole moments (TDM in D) of the first two electronic transitions of CAN calculated using different geometries and environments.

| | $S_0-S_1$ ($2A_g^-$) | | $S_0-S_2$ ($1B_u^+$) | |
| --- | --- | --- | --- | --- |
| | $\Delta E$ | TDM | $\Delta E$ | TDM |
| CAN in gas phase (gas-phase geometry) | 2.74 | 0.0 | 2.85 | 17.4 |
| CAN in gas phase (OCP-PBS geometry) | 2.13 | 0.3 | 2.57 | 17.4 |
| CAN in OCP-PBS (MD average) | 2.00 | 7.8<sup>a</sup> | 2.23 | 19.0<sup>a</sup> |
| CAN in gas phase (OCP<sup>O</sup> geometry) | 2.28 | 0.6 | 2.64 | 18.6 |
| CAN in OCP<sup>O</sup> (MD average) | 2.24 | 3.3<sup>a</sup> | 2.53 | 18.4<sup>a</sup> |

<sup>a</sup> The TDM average was obtained as root-medium-square of the TDM vectors calculated along the restMD trajectories

To assess the impact of the flexible linker on the properties of CAN, we performed an additional MD simulation excluding the OCP linker residues G169-P179. In this model the calculated S_0_-S_1_ TDMs are slightly smaller, while the corresponding transition energies do not differ significantly from the ones with the linker (Fig. S7). Therefore, the flexibility of the linker only has a minor effect on the optical properties of CAN.

To quantitatively assess the importance of a well resolved structure around the CAN we repeated the same restMD and QM/MM calculations on the OCP-PBS complex, utilizing our previous structure restricted to a resolution of 2.6 Å (*4*). This analysis allowed a direct comparison of the dark state properties (Figure 3d-f). The previous structure gave significantly smaller TDMs and higher excitation energies than the structure from our present work. Specifically, our earlier structure of OCP-PBS did not give significantly different TDMs relative to OCP^O^, whereas the TDMs obtained on the present structure are much larger (Figure 3d).

The comparison with OCP^O^ also shows that the dark state energy is red shifted by ∼0.2 eV in OCP-PBS, with a somewhat broader distribution. Furthermore, TDMs in OCP-PBS are substantially larger than in OCP^O^. The average value of S_0_-S_1_ TDM in OCP^O^ is around 3 D and its distribution is centered between 1 and 2 D, which is close to the earlier calculations (*21*). This comparison suggests that the specific electrostatic environment of CAN in OCP-PBS substantially increases the TDM of the S_0_-S_1_ transition. In fact, looking at results calculated using a geometry of CAN optimized in gas-phase and the ones imposed by the OCP-OBS complex and the OCP^O^, respectively (Table 1), it is clear that the differences in the properties of CAN cannot be explained only in terms of a geometrical effect. A similar behavior is found for the overall absorption spectrum of CAN in OCP-PBS and OCP^O^ (Figure S10), in agreement with experiments (*22, 23*).

The analysis of QM/MM simulations shows that the significant TDM of the dark state of CAN in OCP-PBS is mainly induced by a marked imbalance of charge distribution within the CAN binding pocket. By analyzing the effect of each residue separately (see Methods), we found that almost all charged residues close to CAN, in OCP or in the PBS, contribute to the increase of the TDM (Fig 4). Proximal to the PBS, R155 in OCP has the largest effect on the TDM, followed by some positively charged residues on ApcA (K61, K62) and ApcB (K53). At the same time, the negatively charged residues, E34 and D35, which are located on the distal, solvent-exposed surface of OCP, also increase the TDM. The only charged residue that decreases the TDM is D64 on ApcA, which is needed to make a salt bridge with R62. Thus, it appears that the positive and negative residues on either end of the CAN give rise to an increase in the S_0_-S_1_ TDM by generating a strong electrostatic potential difference that breaks the symmetry along the conjugated chain of the carotenoid. Several highly conserved residues of the PBS located at the OCP binding interface (ApcA-K61, ApcA-R62, and ApcB-K53) contribute to the TDM increase, suggesting that OCP-PBS binding is essential to achieve a large TDM in the CAN. Notably, removing any of the charged residues at either end of the CAN results in at least threefold decrease of the TDM. Although this estimate only covers the direct electrostatic effect of the residues (See Methods), this analysis strongly suggests that only in the OCP-PBS complex the dark state of CAN can acquire a large TDM.

**Figure 4:**
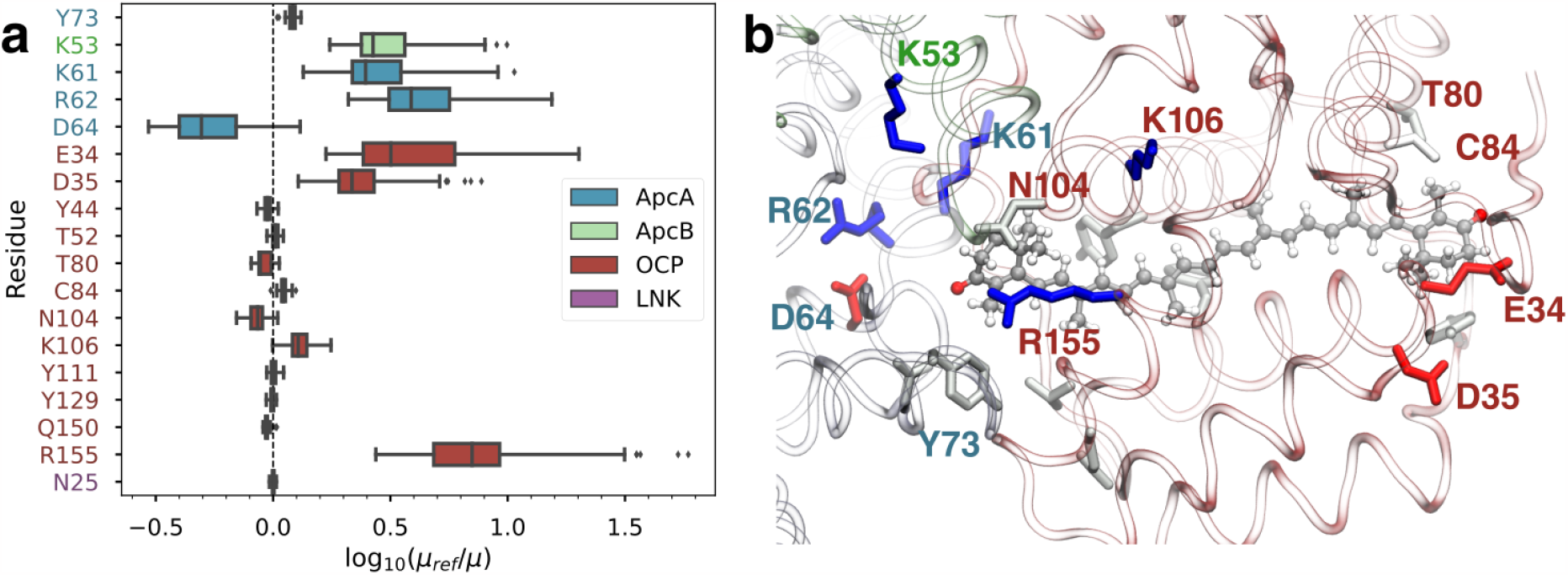
Impact of the molecular environment on the CAN S_1_ TDM. (**a**) Effect of selected residues on the TDM of CAN (S_1_ state), computed as the logarithm of the ratio between the full OCP-PBS calculation (*μ*^ref^) and the calculation excluding the selected side chain (*μ*). (b) Representation of the most important residues in the CAN binding pocket. Colors indicate the residue type: blue for Lys/Arg, red for Asp/Glu, and grey/white for polar/nonpolar residues. Label colors refer to the sidechain with the same color code as in (a).

Our calculations indicate that the distortion of the carotenoid in OCP-PBS is not sufficient to achieve a significant TDM for the S_0_-S_1_ transition. Instead, the charges of the protein environment substantially increase the S_0_-S_1_ TDM by creating an electrostatic gradient along the CAN conjugated chain. The TDM increases by virtue of a mixing between the pure 2A_g_ and 1B_u_ states: in fact, the S_1_ state borrows dipole strength from the S_2_ state (Fig. S5). Our calculations show that the S_1_ and S_2_ states are close in energy, at least at the Franck-Condon point (Fig. 3b, Fig. S5), and the increase in S_0_-S_1_ TDM is only substantial when there is a small energy difference between S_1_ and S_2_ (Fig. S5). We finally note that even if such an increased TDM (∼8 D) should be visible in the absorption spectrum, our simulations shows that the corresponding band is hidden in the red edge of the more intense S_0_-S_2_ band (Fig. S5d).

### Energy transfer and OCP-mediated quenching

The same QM/MM calculations along the restMD trajectories were used to compute excitation energy transfer (EET) couplings of CAN with the closest PCB pigments. The closest PCBs are those of two different ApcA subunits; one of them is only 10-15 Å away from the CAN (Figure 5a). Therefore, the point-dipole approximation is not fully justified here, and we employed the TrEsp method (*24*) for calculating the CAN(S_1_)-PCB couplings (see Methods in the Supplementary Materials). The coupling distributions (Figure 5b) are quite broad and reflect the observed variability in the TDM. Indeed, both CAN-ApcA couplings are essentially proportional to the TDM of CAN (Figure S8). This result is expected because both the TrEsp charges and the TDM are proportional to the mixing between the states. The mean couplings, 54 cm^−1^ for ApcA_1_ and 27 cm^−1^ for ApcA_2_, are compatible with significant PCB-to-CAN EET rates. The other (ApcB) pigments have couplings smaller than 8 cm^−1^.

**Figure 5:**
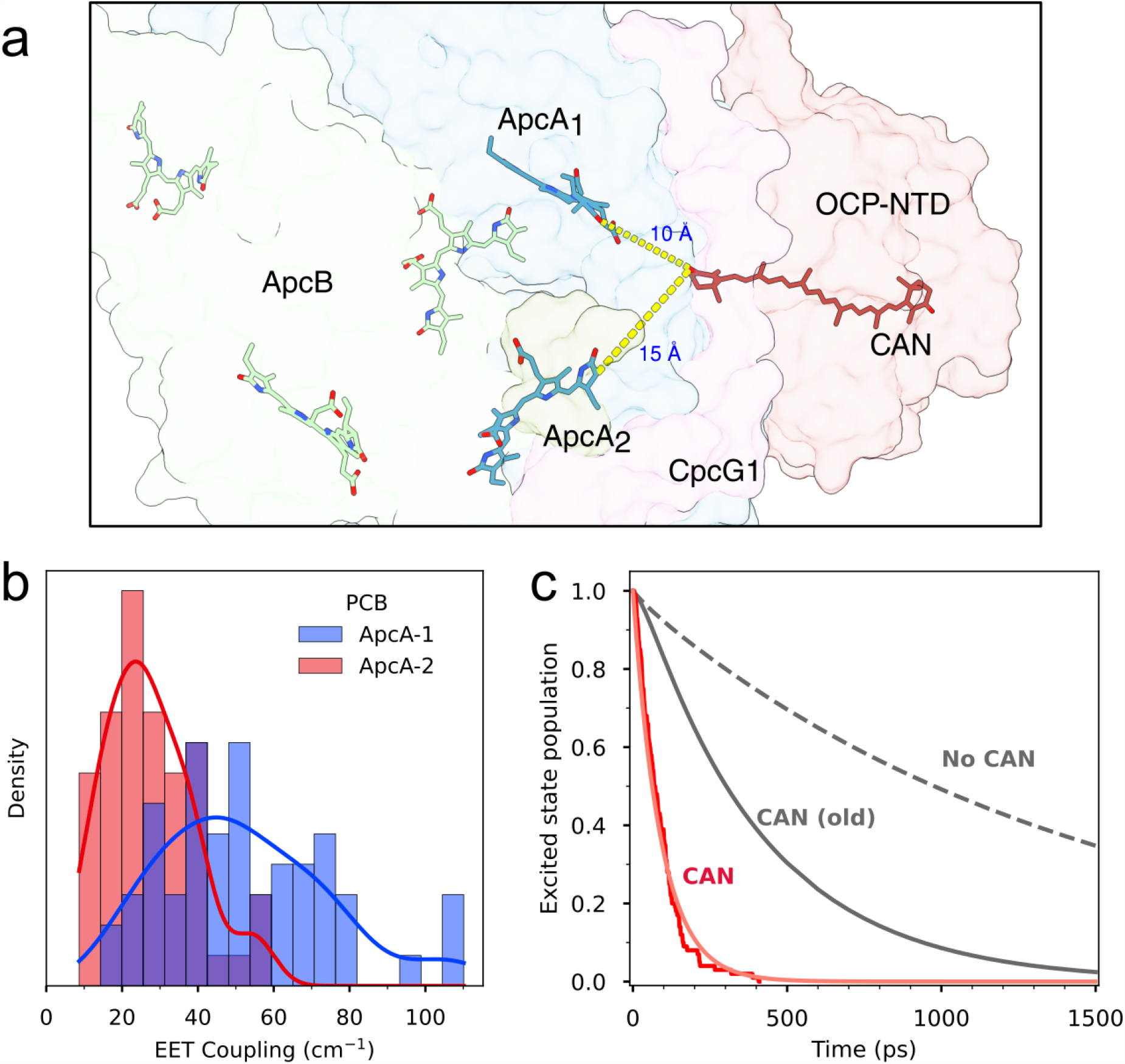
Simulation of EET quenching and the excited-state decay in the OCP-PBS complex. (a) Representation of CAN in OCP and the closest Apc pigments. The closest ApcA pigments are denoted as ApcA_1_ and ApcA_2_. (b) Distribution of the calculated EET couplings between CAN S_0_-S_1_ and ApcA_1/2_. (**c)** Results of 100 stochastic simulations of PBS decay in the presence of four CAN parametrized with QM/MM simulation results (red), excited state decay simulation from ref. (*4*) for OCP-PBS quenched by four CAN (grey solid line), unquenched PBS (dashed grey line).

We employed the CAN(S_1_)/PCB couplings to calculate the quenching of PBS by CAN in OCP. Simulation of the excitation energy transfer dynamics was performed essentially as described in Dominguez-Martin et al. (*4*). Here, however, the point-dipole approximation was used only for the PCB-PCB couplings within PBS to model the energy flow through PBS, whereas the individual couplings between CAN and the two nearest ApcA PCBs were directly taken from the QM/MM TrEsp calculations. The other CAN-PCB couplings were neglected, because our previous analysis showed their contribution to the quenching is negligible (*4*). The CAN-PCB(ApcA) couplings as well as the S_0_-S_1_ energies of CAN were randomly sampled from the QM/MM distributions shown in Fig 5b and Fig 3b, and assigned independently to each of the four OCP carotenoids. We have used the same estimated S_0_-S_1_ absorption band shape used in (*4*), but with distribution of absorption maximum according to Fig 3b. The simulation was then run using the stochastic approach, starting from a randomly selected rod phycocyanobilin. For each run the CAN parameters were independently sampled. The results of simulations are shown in Fig 5c. The effect of the increased CAN-PCB coupling due to large S_0_-S_1_ TDM of CAN in OCP-PBS is evident. The overall PBS lifetime drops to ∼100 ps providing all four OCPs are attached to PBS, in agreement with a lifetime of ∼160 ps which has been determined experimentally (*17*).

## Discussion

We have presented the high-resolution cryo-EM structure of a cyanobacterial phycobilisome in complex with OCP and characterized its inherent flexibility.

Some degree of flexibility is inherent in all protein complexes, but how flexibility impacts protein function is only beginning to be understood. The stochastic sampling of small conformational changes can immediately translate to changes in protein activity, as has been shown for HIV protease or triose-phosphate isomerase (*25–28*). Ensemble averaging techniques like NMR or computational modeling can be used to describe such motions, but these methods are still limited in the size of the system that can be investigated. On the other hand, a common drawback of cryo-EM structural studies is that they suffer from insufficient resolution. Hence, only a few complex dynamic systems, such as the translating ribosome, have been successfully investigated in detail (*29–31*). Continued progress in cryo-EM sample preparation and data processing holds unique promise for future studies of dynamic protein complexes, by improving the achievable resolution and allowing more sophisticated analysis of the inherent motion within the dataset.

In examining the OCP-PBS complex, we identified several movements with potential implications for its light-harvesting function. In particular, the inherent movement of the PBS rods hints at a conformational switching that controls access of OCP binding to PBS (*4*). Other regions of the PBS, such as the top cylinder and the NTD of the OCP are remarkably stable. In contrast, there is considerable flexibility in the CTD dimer and the NTD-CTD linker, potentially affecting stability or longevity of the quenched PBS. The role of the observed intrinsic motions in regulating light harvesting can now be tested.

The substantially increased accuracy achieved in our atomic model allowed us to perform a detailed QM/MM simulation of EET quenching of PBS by OCP. Our results predict a substantially larger S_0_-S_1_ TDM for CAN in OCP-PBS than previously estimated (*21*). This leads to larger EET couplings of CAN with the closest ApcA phycocyanobilin (Fig. 3d), making the OCP an efficient quencher. The simulation results (Fig. 5c) show that tuning the excited state properties of CAN by the environment in OCP-PBS significantly enhances the quenching rate. While in our previous simulation (*4*), using parameters estimated from other systems, we obtained a 300-400 ps time constant for APC-to-CAN energy transfer, by using the QM/MM calculations on the real system, the quenching time decreases to ∼100 ps (Fig. 5c), which corresponds well to the value obtained from time-resolved fluorescence experiments (*17*). This clearly demonstrates the importance of CAN-protein interactions in tuning the quenching, since they enhance the quenching rate nearly four fold compared to that calculated earlier without involving the effect of these interactions (Fig. 5c). The variability observed along our restMD trajectories in EET couplings (and consequently in EET rates) not only suggests that the efficiency of EET quenching is highly tunable by the protein environment, but also explains the heterogeneity observed in quenched PBS (*32*).

Comparing our simulations on the high resolution structure with the previous one, we showed that the 2.6 Å resolution in the previous structure could not provide qualitatively correct results in the QM/MM calculations. Given the extreme sensitivity of the CAN electronic structure to the electrostatic environment (Figure S11), even small differences in the structure could reflect dramatically on the QM/MM results. In particular, we can observe a different position of R155 within the OCP NTD, which is located close to the CAN end and has a substantial effect on its excited-state properties (Figure 4). Furthermore, other differences can be found in the charged residues of ApcA in the PBS. Overall, the improved structural resolution allowed us to better pinpoint the position of charged side chains, which ultimately determines the success of our QM/MM strategy.

The calculations using the high resolution structure confirm that to achieve an efficient quenching, there is no need to quench the lowest energy state emitting at 680 nm, which is associated with ApcD and ApcE subunits (*33, 34*). Instead, quenching of ApcA pigments emitting at 660 nm is enough to provide photoprotection, as suggested in earlier reports based on analyses of time resolved fluorescence data (*35, 36*). Moreover, interaction of OCP with ApcA provides a natural way for switching between the quenched and non-quenched state of the PBS. First, changing the position of the rods can prevent OCP binding by blocking the binding site (*4*), thus keeping PBS in an unquenched state. Second, when OCP is bound, the binding site can still allow OCP to interact with FRP, which is needed to revert to inactive OCP^O^. Such regulation would be complicated (if not impossible) to achieve if the quenching site were associated with the lowest energy state, because of the limited access to the ApcE and ApcD subunits (*4*). In summary, our study illuminates structural dynamics and energy transfer quenching processes within the cyanobacterial phycobilisome through high-resolution cryo-EM and multiscale quantum chemical calculations. These details open new avenues for fundamental research on energy transfer in pigment-protein complexes. Likewise, this high resolution picture of the PBS and its mechanism of NPQ serve as inspiration for synthetic biologists, chemists and materials scientists to design new sustainable technologies for harnessing the clean and abundant energy in sunlight.

## Acknowledgments

We thank Abhiram Chintangal and Kurt Stine for computational support. We thank Bong-Gyoon Han and Robert Glaeser for help with the streptavidin affinity grid technique. The research in the Kerfeld lab was supported by the Office of Science of the U.S. Department of Energy under award number DE-SC0020606. EN is funded by the National Institutes of Health, NIGMS grant GM127018. She is a Howard Hughes Medical Institute Investigator. L.C., M.B. and B.M. acknowledge funding by the European Research Council under the Grant ERC-AdG 786714 (LIFETimeS). D.B. and T.P. thanks the Czech Science Foundation (19-28323X) for financial support.

## Author contributions

PVS performed cryo-EM analysis and data interpretation. LC performed QM/MM and restMD calculations, interpreted data. MS built the atomic model, interpreted data. MB helped perform QM/MM and restMD calculations, interpreted data. MADM and HK prepared the sample. DB performed EET quenching simulations and interpreted data. AK and AFK performed cryo-EM data collection and provided expertise. BJG, EN, TP, BM and CAK provided expertise. BM, CAK and EN supervised the project. PVS and LC wrote the manuscript with input from all authors.

## Data availability

The atomic coordinates have been deposited in the PDB with the accession codes 8TPJ for the T-cylinder disk bound to OCP, 8TO2 for the B-cylinder bound to OCP, 8TRO for the rod and 8TO5 for the central rod disk in C1 symmetry. The electron microscopy maps have been deposited in the Electron Microscopy Data Bank with the accession codes 41463 for the holo OCP-PBS complex, 41475 for the T-cylinder disk bound to OCP, 41434 for the B-cylinder bound to OCP, 41585 for the rod, 41435 for the central rod disk in C1 symmetry, and 41436 for the central rod disk in D3 symmetry. The raw micrographs for all datasets have been deposited in the Electron Microscopy Public Image Archive with the accession code EMPIAR-11644. The code used in this study to subtract the streptavidin lattice from the electron micrographs is available on GitHub at https://github.com/pvsauer/StreptavidinLatticeSubtraction.

## Supplementary Materials

Methods

Figs. S1 to S11

Table S1

References

Movies S1 to S3

