## Supplementary Materials for "Structural and quantum chemical basis for OCP-mediated quenching of phycobilisomes"

Methods
Figs. S1 to S11
Table S1
Captions for Movies S1 to S3
References (37–52)

### Methods

#### High resolution cryo-EM data acquisition and processing

Plunge frozen grids prepared and imaged previously *(4)*were provided to Thermo Fisher Scientific, Eindhoven, Netherlands for data collection. A total of 16,284 exposures were collected on a Krios G3i microscope operated at 300kV and equipped with a cold field emission electron gun, Selectris X energy filter and Falcon 4 camera. Exposures were collected with a nominal pixel size of 0.727 Å and recorded in EER file format with 1176 total raw frames per movie using aberration free image shifting and fringe-free imaging *(5,37)*.The defocus range was between -0.4 and -1.2 micrometers and the total dose per exposure was 40 electrons/$Å^2$.

Images were motion corrected using motioncor2 with variable frame grouping optimized for early frames with high beam induced motion *(38)*.During motion correction, frames were temporarily upscaled from 4K to 8K to improve signal-to-noise at high resolutions. After motion correction, the streptavidin lattice was subtracted using in-house scripts *(9,10)*.For subsequent processing motion corrected and subtracted micrographs were imported into Cryosparc v.4.2 *(9,10,39)*.After CTF estimation, PBS particles were picked using our previous structure as a template. After several rounds of 2D and 3D classification, as well as global and local CTF refinement, we obtained a single class corresponding to OCP-PBS at a global resolution of 2.2 Å according to the FSC = 0.143 criterion *(40)*. To analyze flexibility of the holo OCP-PBS we performed 3D variability analysis as implemented in Cryosparc, filtering the map to 20 Å and using 3 modes *(12)*.

To obtain a better map of the central core of OCP-PBS, in lieu of local refinements, particles centered on coordinates for holo complexes were re-extracted at smaller box sizes, effectively truncating the peripheral rods. The map was refined to 2.2 Å and used for subsequent flexibility analysis and local refinements. For flexibility analysis, we ran 3D variability analysis with a 4 Å filter and 3 modes *(13)*.

After symmetry expansion, local refinement of the T-disk yielded a 2.1 Å map, which was used as a basis for 3DFlex analysis as implemented in Cryosparc, which we ran using 5 latent dimensions and 307,000 particles.

To obtain reconstructions of the rods, templates from our previous structure were used to pick and extract rod particles in a separate workflow from holo-PBS particles. After 2D classification, 3D classification, global and local CTF refinement we obtained a final map at 1.9 Å resolution. Local refinement of the central rod disk yielded a reconstruction at 1.8 Å.Applying D3 symmetry then yielded a reconstruction at 1.6 Å, though it lacked the asymmetric central linker protein portion.

#### Atomic modeling, hydrogen modeling

Atomic models were built by first fitting the respective existing structures or subsets of them (pdb IDs 7SCC, 7SCB, 7SCA) into the cryo-EM density using UCSF ChimeraX 1.3 *(41)*.Hydrogen atoms were added using phenix.reduce *(42)*and a morph step was performed during the initial refinement with phenix.refine 1.19.2 to fit the model into the density. Manual rebuilding with COOT 0.97 *(43)*was alternated with refinement with phenix.refine 1.19.2. Water molecules were added for the final rounds of refinement using phenix.douse and manually evaluated by inspection in COOT 0.97. In order too visualize the signal from hydrogen atoms in the 1.6 and 1.8 Å maps of the central rod disk, we used the program Servalcat as implemented in Refmac5 and ccpEM v. 1.6.0. to refine the atomic model and create Fo - Fc maps *(44)*.

Structures were visualized with UCSF ChimeraX *(41)*

#### Molecular Dynamics

System preparation and all MD simulations were performed using Amber 18 *(45)*.The model system (OCP-NTD and the seven closest apoprotein chains of PBS core) was solvated in a truncated octahedron box of $\sim$14 nm diameter, ensuring at least 4 nm spacing between periodic images of the protein complex. $Na^+$ ions were added to neutralize the system. The protein was described with the AMBER ff14SB force field *(46)*,whereas for CAN we used our previously developed force field *(19)*,and GAFF for PCBs *(47)*.Water was described with the TIP3P model. The entire system was minimized subject to 4 kcal mol$^{-1}$ Å$^{-1}$ restraints on all non-solvent non-hydrogen atoms. Then, the system was heated gradually to 300 K in 20 ps, with the same restraints in the NVT ensemble. Finally, the box was equilibrated through a 1 ns NPT simulation using the Monte Carlo barostat implemented in Amber, keeping the restraints. All simulations were run with the Langevin thermostat, a time step of 2 fs, and the SHAKE algorithm. Periodic PME electrostatics was used with a short-range cutoff of 1 nm. The restMD production was run for 40 ns subject to 4 kcal mol$^{-1}$ Å$^{-1}$ restraints on the backbone of the proteins. Only the OCP linker residues (169-179) were allowed to move freely. The CAN molecule was frozen to the previously QM/MM optimized geometry. For the geometry optimization of CAN we used Density Functional theory (DFT) at the B3LYP/6-31G(d) level.

#### Excited-state QM/MM calculations

Excited states of CAN were determined with a semiempirical CI method (SECI) with parameters optimized specifically for carotenoids *(20)*.Excited-state calculations on the bilins were performed using time-dependent Density Functional theory (TD-DFT) at the CAM-B3LYP/6-31G(d) level. For calculations in OCP-PBS, we used an electrostatic embedding QM/MM scheme including point charges of the protein, other cofactors, water molecules, and ions. Transition charges were obtained from the transition electrostatic potential, as obtained from QM/MM calculations, in accordance with the TrEsp method *(24)*.The coupling between pigments A and B was calculated as

$$V_{\mathrm{AB}} = \sum_{i}^{A} \sum_{j}^{B} \frac{q_i q_j}{r_{ij}}$$

where $q_i$ and $q_j$ are the transition charges on atoms $i$ of pigment A and atom $j$ of pigment B respectively, and $r_{ij}$ is the distance between them.

The effect of protein residues on the excitation energy and TDM of CAN's excited states was determined as follows: for each considered residue, we repeated all the QM/MM calculations after setting to zero all the charges of its side chain. The resulting TDM ($\mu$) was then compared to the reference QM/MM calculation that includes all charges ($\mu^{\mathrm{ref}}$), and the logarithm of the fold change $\log(\mu/\mu^{\mathrm{ref}})$ was calculated for each snapshot of the MD. Note that this analysis neglects any structural changes that may arise from removing the residue.

#### Energy transfer modeling

Simulation of the excitation energy transfer dynamics has been performed essentially as described in Ref.*(4)*.We employed Förster theory to compute the pairwise energy transfer rates, from couplings ($V$) and overlaps ($J$) of normalized absorption and emission spectra, using equation $k_{AB} = 1.18\ V_{AB}^2\ J$ *(48)*.As the full absorption profile of the CAN $S_0$-$S_1$ transition is not known, we employed the emission lineshape of another ketocarotenoid, peridinin *(49)*,assuming a mirror image relationship between absorption and emission lineshapes *(50)*A detailed balance condition was applied to account for uphill energy transfer rates. The simulation was then run using the stochastic approach, using the Gillespie algorithm *(51,52)*.Each simulation run started from a randomly selected rod PC bilin and for each run the carotenoid parameters were independently sampled.

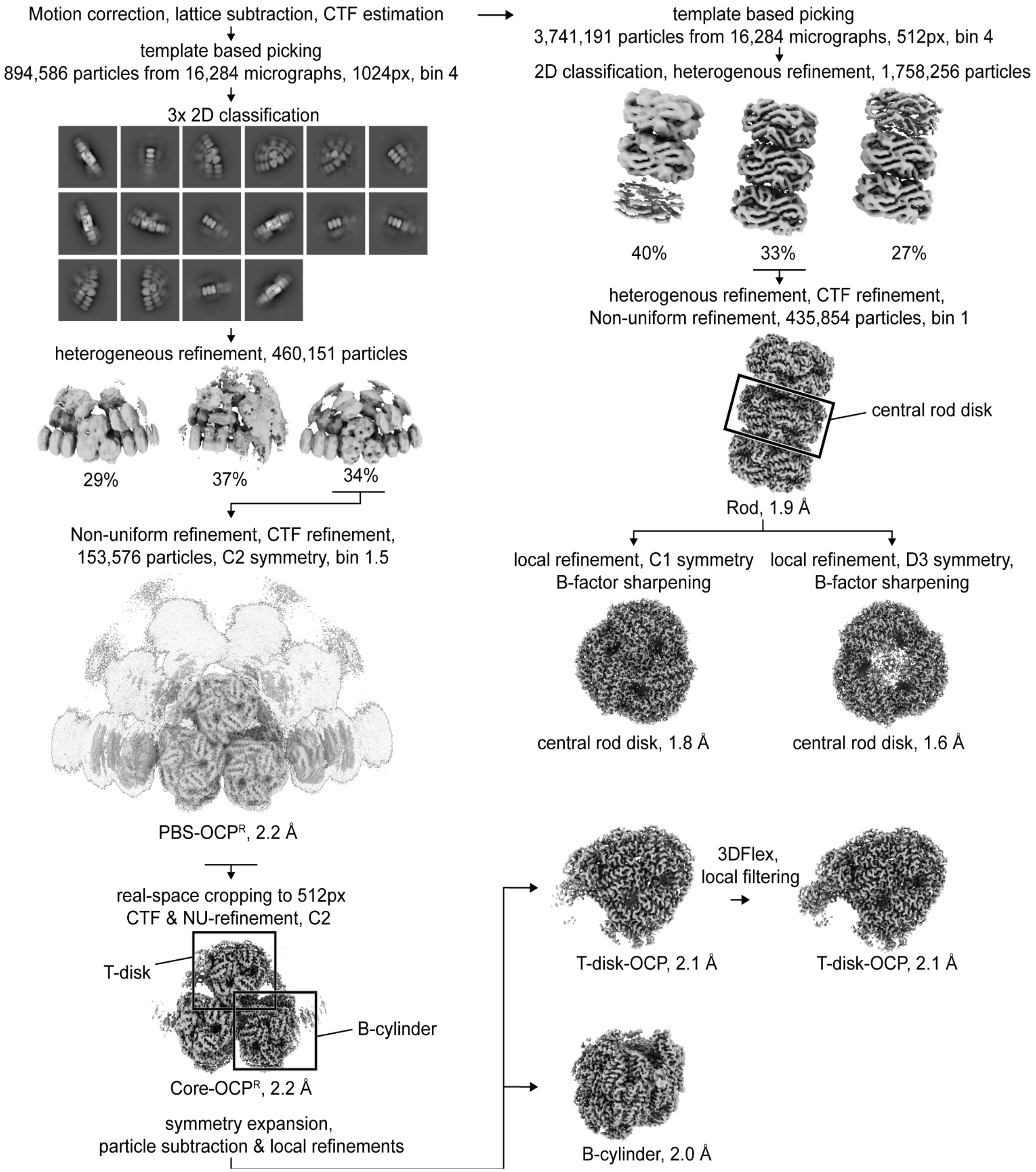


Figure S1: Cryo-EM processing workflow. Holo-OCP-PBS particles and the rods were processed independently.

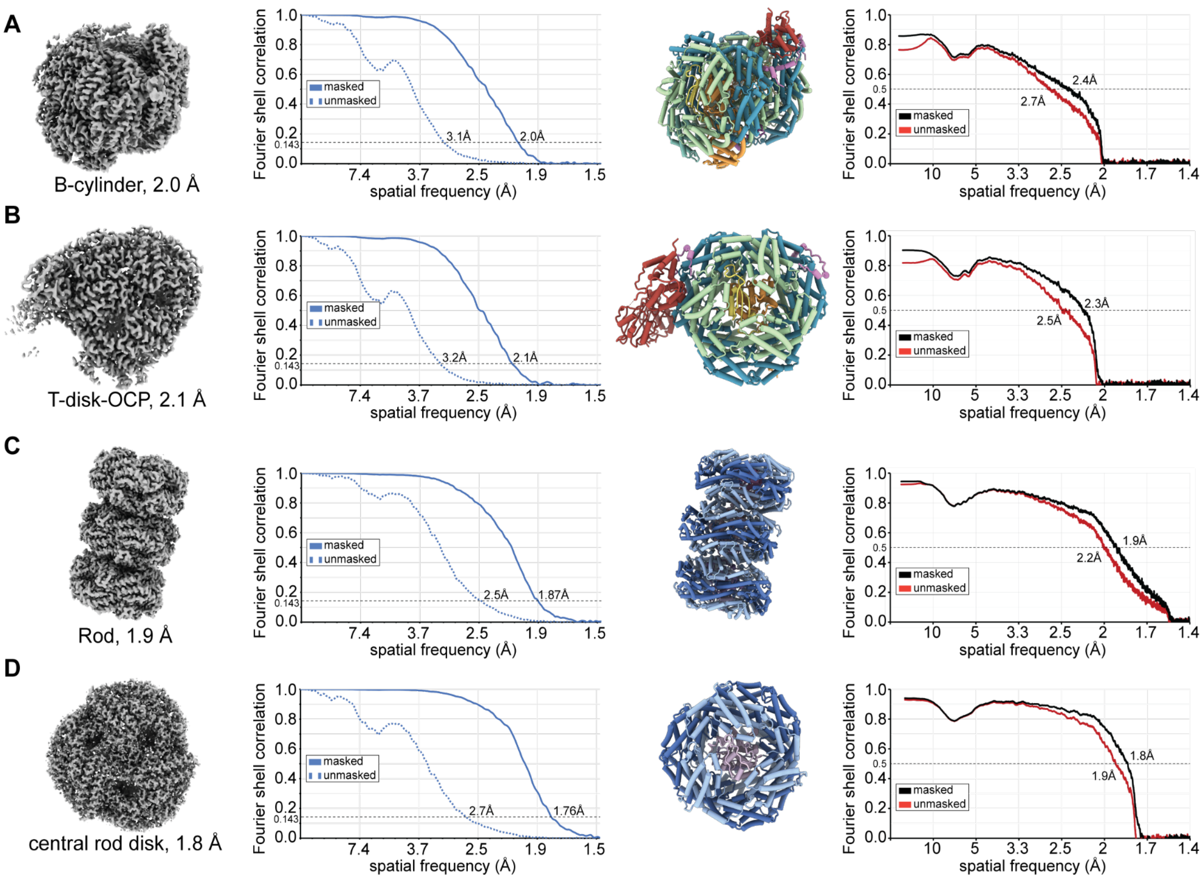


Figure S2: Fourier shell correlation curves and model validation. From left to right, final cryo-EM reconstructions (identical to Figure S1), their corresponding Fourier shell correlation curves, their atomic models and the map-to-model validation plots, shown for the B-cylinder-OCP (A), the T-disk-OCP (B), the rod (C), and the central rod disk (D). The entire OCP-PBS model can be put together from the partial models (A)-(C) and the cryo-EM map of the complete complex.

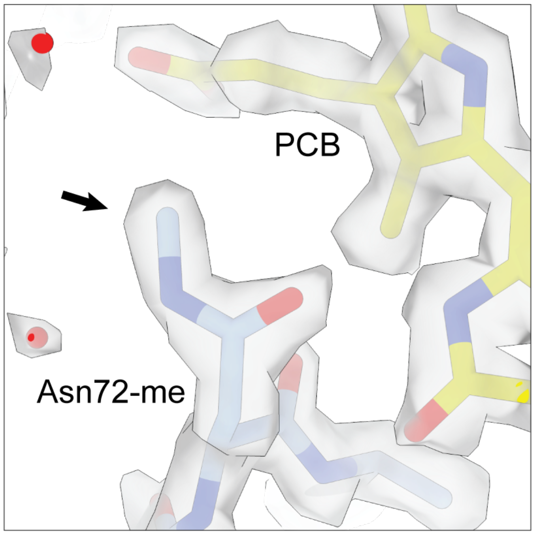


Figure S3: N4-methylasparagine in position 72 of CpcB, presumably the result of post translational modification, adjacent to a phycocyanobilin (PCB). Arrow indicates additional methyl group.

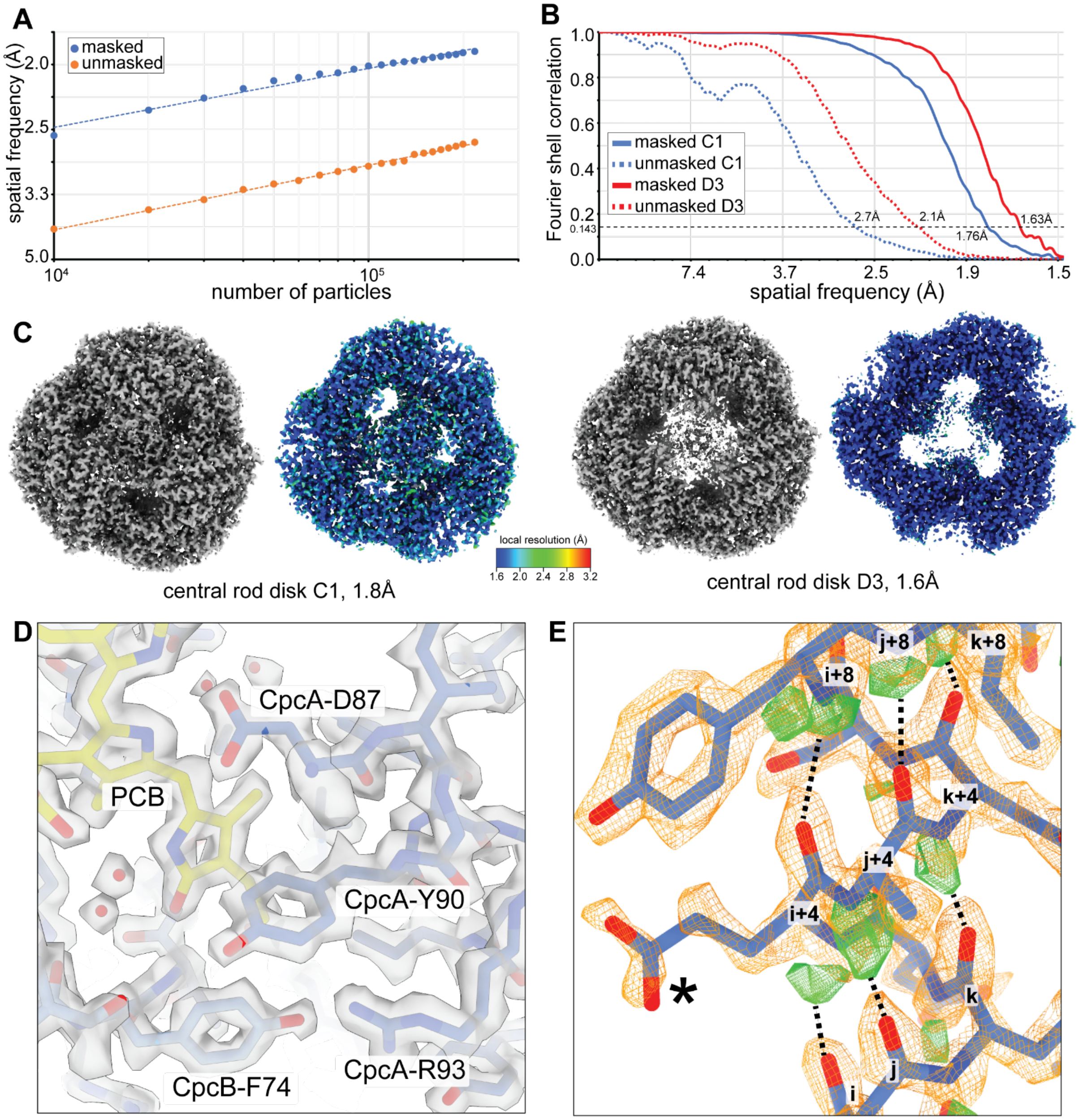


Figure S4: High resolution details of the central rod segment. (A) ResLog plot of the full rod showing that attainable resolution is limited by particle numbers. (B) Fourier shell correlation curves from local reconstructions of the central rod disk, using either C1 or D3 symmetry. (C) Cryo-EM maps of the central rod disk either using C1 symmetry (left) or D3 symmetry (right). Next to each map a slice of the local resolution map is presented, showing the resolution in the interior. (D) High resolution map detail of the 1.6 Å D3 symmetry rod disk map. Water molecules and holes in aromatic residues are clearly visible. (E) FO map of an alpha helix from the C1 symmetrical central rod disk (orange mesh). The FO-FC map (green mesh) shows differential density indicating the potential position of hydrogen atoms. The hydrogen positions are in agreement with the hydrogen bonding patterns in alpha helices, which bridge the carbonyl moiety of residue i with the peptide bond of residue i+4. Potential hydrogen bonds are indicated with dotted lines. Asterisk marks a terminal carboxyl group of a glutamate residue that has disconnected density from the backbone, presumably a result of beam induced decarboxylation.

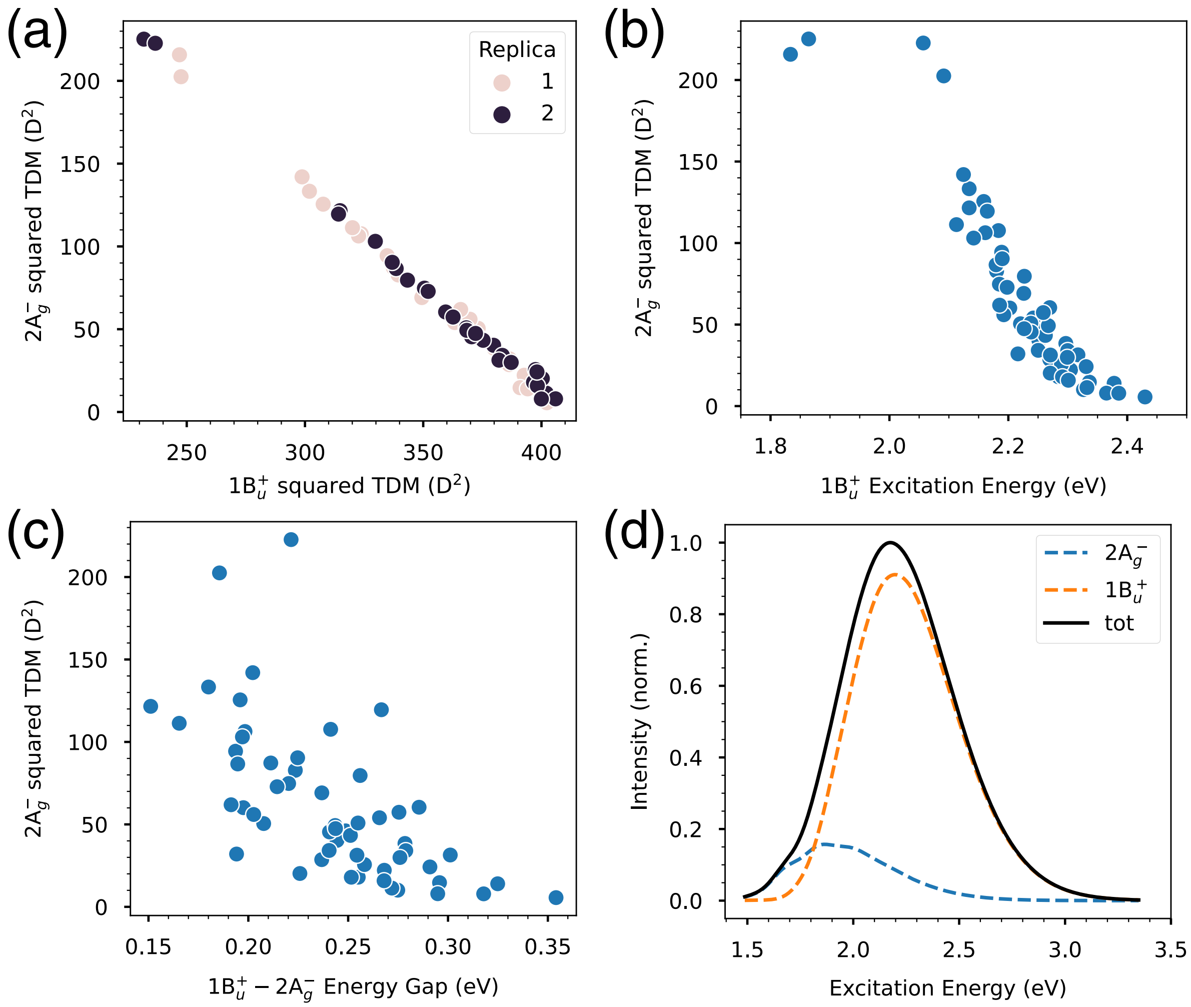


Figure S5: Demonstration of S1-S2 mixing for CAN in OCP-PBS. (a) Correlation between the squared TDM of the $S_0$-$S_2$ ($1B_u{}^+$) and of the $S_0$-$S_1$ ($2A_g{}^-$) transitions. Points of different colors represent structures extracted from two different MD replicas (b) Correlation between the $S_0$-$S_2$ excitation energy and the squared $S_0$-$S_1$ TDM. (c) correlation between the energy difference ($\Delta$E S2-S1) and the squared $S_0$-$S_1$ TDM. (d) Spectrum calculated by convoluting the S1 and S2 contributions with the vibronic lineshape of the two transitions.

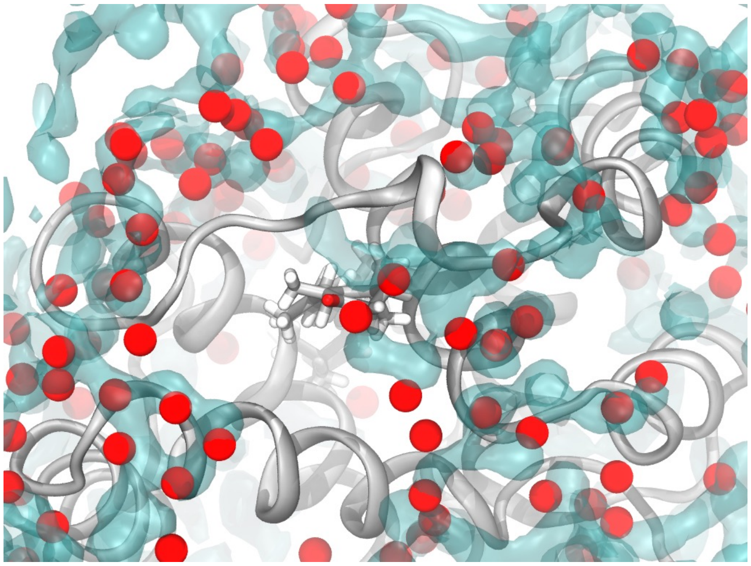

Figure S6: Water occupancy at the OCP-PBS interface during the MD simulations. The OCP NTD is shown (cartoon representation) along with CAN (licorice). The red spheres represent water molecules determined in the Cryo-EM model, whereas the cyan surface shows the regions of high number density ($> 0.04/\text{Å}^3$) of water oxygen atoms. The water density was obtained throughout the restMD trajectories of OCP-PBS

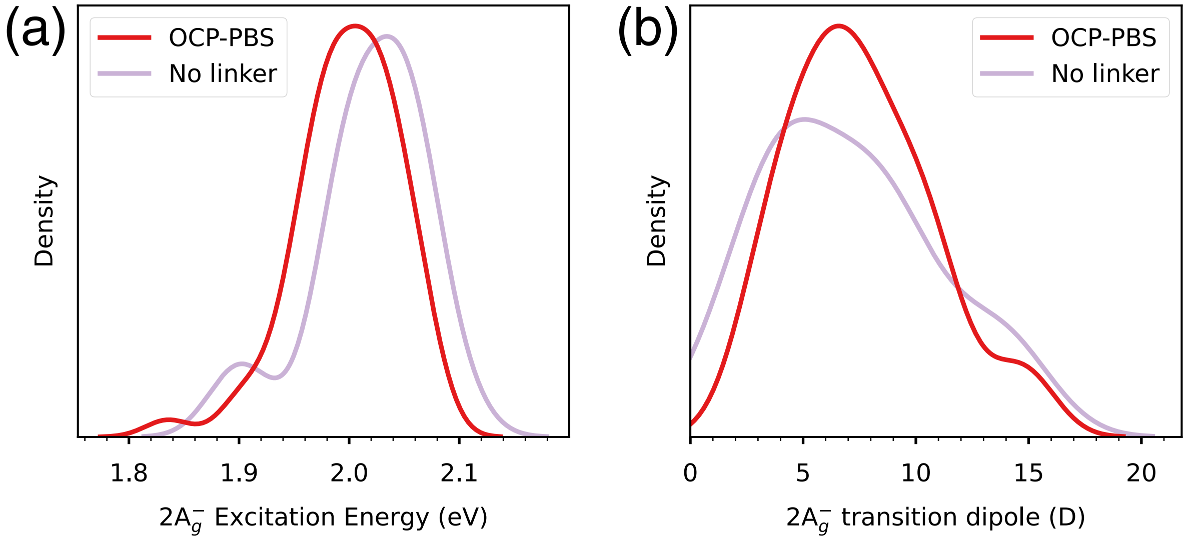


Figure S7: Effect of the linker. Distributions of (a) $S_0$-$S_1$ ($2A_g^-$) transition energies and (b) $S_0$-$S_1$ ($2A_g^-$) transition dipole moments (right) in the OCP-PBS model and without the linker. Data correspond to two pairs of completely independent MD replicas.

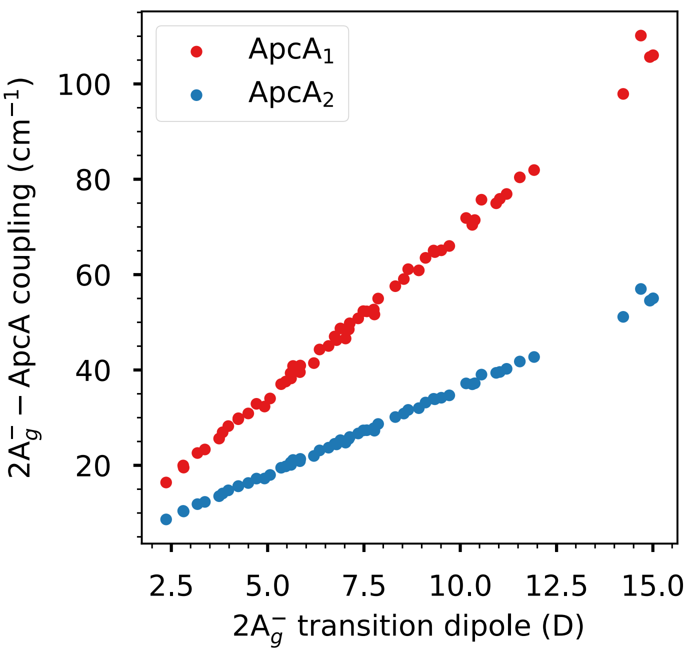


Figure S8: Correlation between the $S_0$-$S_1$ ($2A_g^-$) transition dipole of CAN and the coupling to ApcA1/2.

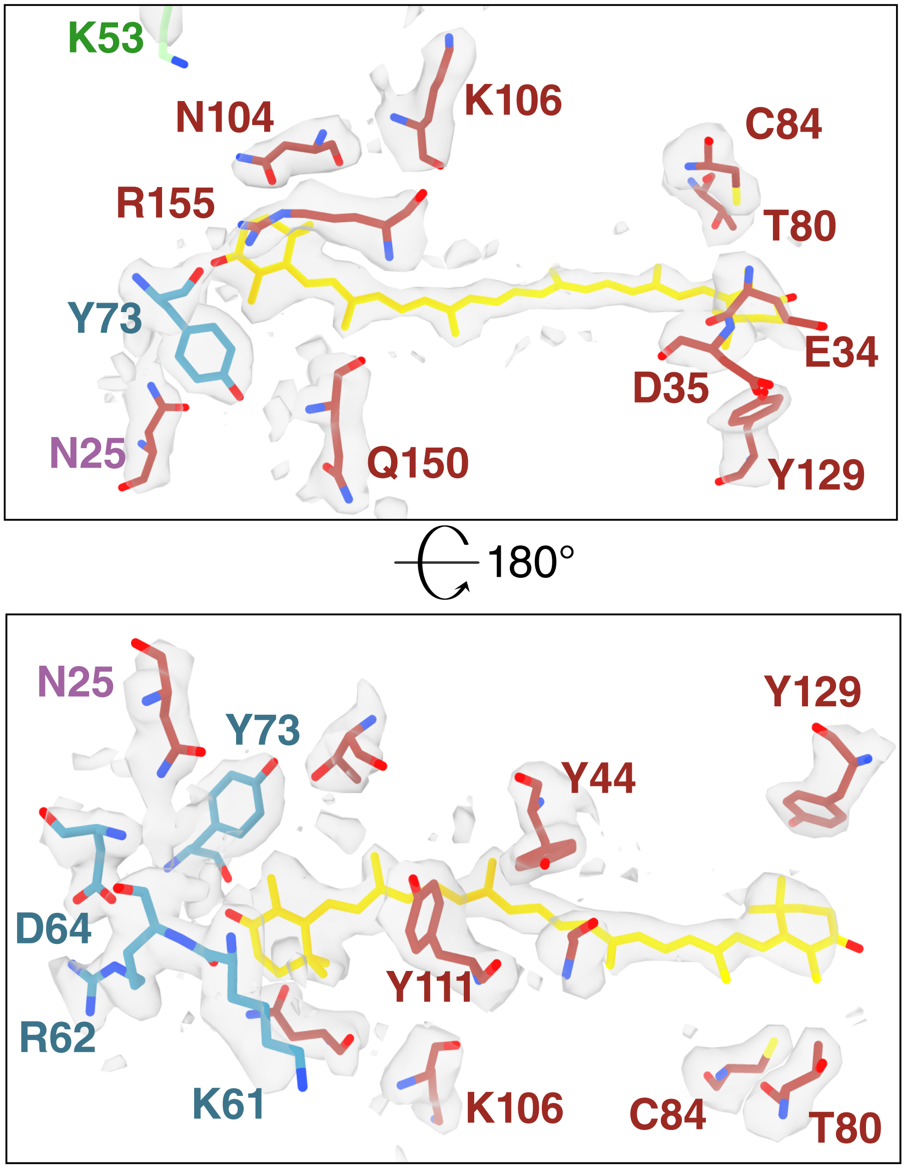


Figure S9: OCP and PBS residues surrounding the CAN in the average MD structure, superimposed to the experimental cryo-EM map. The closest polar or charged residues to CAN are shown.

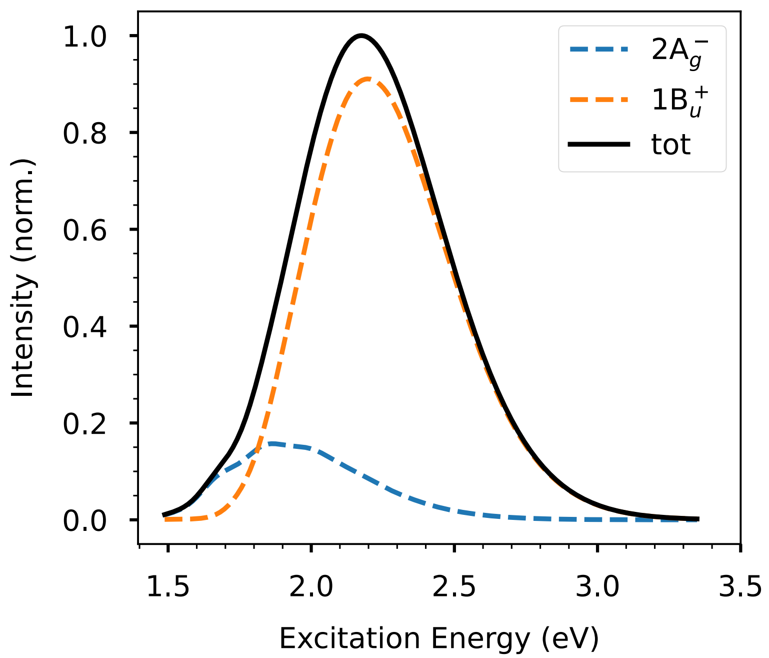


Figure S10: Computed absorption spectra of OCPO and OCP-PBS. The contribution from the S1 state is completely negligible for $OCP^O$, whereas for OCP-PBS the two contributions are shown in Figure S5d.

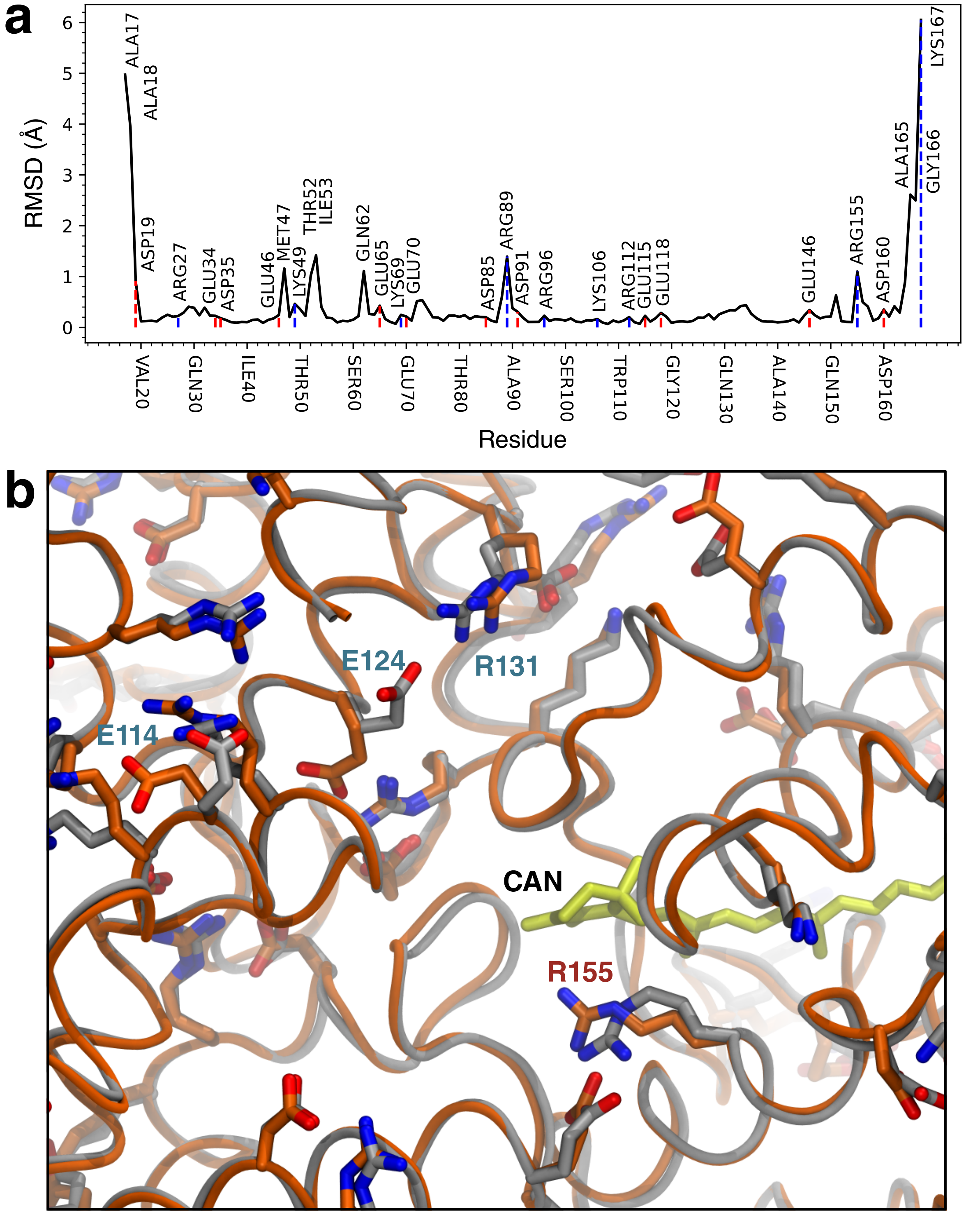


Figure S11: Differences between previous and current structure. (a) Per-residue minimum RMSD between frames of the restMD simulations on the two structures. Charged residues and residues with more than 1 Å RMSD are labeled. (b) Differences between representative restMD samples of the low resolution (grey) and high resolution (orange) structures at the OCP-PBS interface. Only charged residues are shown, and residues showing significant differences are labeled (red labels refer to the OCP sequence, whereas cyan labels refer to the ApcA sequence).

Table S1: Cryo-EM data collection and refinement parameters

| **Title** | **Holo OCP-PBS** | **T-disk OCP** | **B-cylinder** | **rod** | **central rod disk C1** | **central rod disk D3** |
|---|---|---|---|---|---|---|
| **EMD**<br>**PDB** | 41463 | 41475<br>8TPJ | 41434<br>8TO2 | 41585<br>8TRO | 41435<br>8TO5 | 41436 |
| | **Data collection and processing** | | | | | |
| Magnification | 192 000 x | | | | | |
| Voltage (kV) | 300 | | | | | |
| Electron exposure (e–/Å$^2$) | 40 | | | | | |
| Defocus range ($\mu$m) | 0.4 - 1.2 | | | | | |
| Pixel size (Å) | 0.727 | | | | | |
| Symmetry imposed | C2 | | | C1 | | D3 |
| Initial particle images (no.) | 894 586 | | | 3 741 191 | | |
| Final particle images (no.) | 153 576 | | | 435 854 | | |
| Map resolution (Å)<br>FSC threshold 0.143 | 2.2 | 2.1 | 2.0 | 1.9 | 1.87 | 1.63 |
| Map resolution range (Å) | 2-12 | 1.8-12 | 1.8-4 | 1.8 - 2 | 1.7 - 2 | 1.5-1.8 |
| | **Refinement** | | | | | |
| Initial model used (PDB code) | | 7SCC | 7SCB | 7SCA | 7SCA | |
| Model resolution (Å)<br>FSC threshold 0.5 | | 2.2 | 2.4 | 1.9 | 1.8 | |
| | Model composition | | | | | |
| Non-hydrogen atoms<br>Protein residues<br>Ligands | | 39096<br>2463<br>13 | 74379<br>4737<br>25 | 110459<br>6835<br>54 | 37853<br>2295<br>18 | |
| | B factors (Å$^2$) | | | | | |
| Protein<br>Ligand | | 22.9<br>20.4 | 19.3<br>16.0 | 15.3<br>14.0 | 25.5<br>24.6 | |
| | R.m.s. deviations | | | | | |
| Bond lengths (Å) (# $>4\sigma$))<br>Bond angles (°) (# $>4\sigma$)) | | 0.006 (0)<br>0.654 (7) | 0.004 (0)<br>0.644 (19) | 0.005 (0)<br>0.647(4) | 0.003 (0)<br>0.619 (4) | |
| | Validation | | | | | |
| MolProbity score<br>Clashscore<br>Poor rotamers (%) | | 1.20<br>4.22<br>0.00 | 1.31<br>5.71<br>0.00 | 1.21<br>4.26<br>0.38 | 1.04<br>2.56<br>0.23 | |
| | Ramachandran plot | | | | | |
| Favored (%)<br>Allowed (%)<br>Disallowed (%) | | 98.59<br>1.41<br>0.00 | 98.14<br>1.78<br>0.09 | 98.45<br>1.55<br>0.00 | 99.29<br>0.71<br>0.00 | |

**Movie S1**
Representative motion of the holo OCP-PBS complex, determined with 3DVA *(12)*. The rods move 'up' and 'down' relative to the core as rigid bodies. The position of OCP in the complex is indicated. Related to Fig. 2A.

**Movie S2**
Representative motion of the OCP-PBS core, determined with 3DVA *(12)*. The core itself displays a rocking and twisting motion. The ApcA/B hexamer that is absent in a subset of particles is indicated. Continuous flexibility causes the OCP density to appear fragmented. Related to Fig. 2B.

**Movie S3**
Representative motion of the T-disk bound to OCP, determined with 3DVA *(12)*. Relative to the T-disk, the OCP-CTD is flexible. It is connected to the OCP-NTD via a flexible linker. The CAN is in close proximity (indicated). Related to Fig. 2C.
